# Functional and structural maturation of auditory cortex from 2 months to 2 years old

**DOI:** 10.1101/2024.06.05.597426

**Authors:** Yuhan Chen, Heather L. Green, Jeffery I. Berman, Mary E. Putt, Katharina Otten, Kylie L. Mol, Marybeth McNamee, Olivia Allison, Emily S. Kuschner, Mina Kim, Luke Bloy, Song Liu, Tess Yount, Timothy P.L. Roberts, J. Christopher Edgar

## Abstract

In school-age children, the myelination of the auditory radiation thalamocortical pathway is associated with the latency of auditory evoked responses, with the myelination of thalamocortical axons facilitating the rapid propagation of acoustic information. Little is known regarding this auditory system function-structure association in infants and toddlers. The present study tested the hypothesis that maturation of auditory radiation white-matter microstructure (e.g., fractional anisotropy (FA); measured using diffusion-weighted MRI) is associated with the latency of the infant auditory response (P2m measured using magnetoencephalography, MEG) in a cross-sectional (2 to 24 months) as well as longitudinal cohort (2 to 29 months) of typically developing infants and toddlers. In the cross-sectional sample, non-linear maturation of P2m latency and auditory radiation diffusion measures were observed. After removing the variance associated with age in both P2m latency and auditory radiation diffusion measures, auditory radiation still accounted for significant variance in P2m latency. In the longitudinal sample, latency and FA associations could be observed at the level of a single child. Findings provide strong support for a contribution of auditory radiation white matter to rapid cortical auditory encoding processes in infants.

## Introduction

Throughout infancy and childhood, changes to brain architecture allow information to be more quickly and efficiently encoded. As an example, an infant’s day-to-day experience drives an iterative process involving the formation of new synaptic connections followed by neural pruning to create more efficient neural networks [1–3]. Another process associated with the more efficient transfer of sensory neural information is myelination – the wrapping of glial cell membranes around an axon – of the thalamocortical tracts. Specifically, the insulation of axons with myelin from the thalamus (the relay station for all sensory systems except the olfactory system) to the cortex facilitates rapid propagation of sensory information (action potentials) through increased white-matter conduction velocity [4, 5]. Understanding the relationship between functional and structural development of the auditory system in infants is important, as a rapid cortical response to auditory signals is necessary for parsing speech sounds and learning to map meaning onto sounds [6–11].

In infants and young children, the maturation of primary sensory neural circuits can be assessed via electrophysiology (electroencephalography (EEG) or magnetoencephalography (MEG)) by examining the averaged response of electrical (EEG) or magnetic (MEG) cortical pyramidal cell activity to repeated presentations of a sensory stimulus, such as a tone or an image [12–17], measurable noninvasively from sensors outside the head. A feature observed across pediatric EEG/MEG studies is that evoked response latencies decrease as a function of age into adolescence, and thus it takes less time to transmit information from the sense organs (e.g., eyes, ears, skin) to the cortex in older than younger children. For example, the primary visual P1/M1 evoked response [18, 19], the primary auditory P2/P2m evoked response [20–25], and the primary somatosensory N1/M1 evoked response [also known as the N20/M20, 26, 27] appear at earlier latencies in older than younger children.

Magnetic resonance diffusion tensor imaging (DTI) provides insight into the microstructural properties of white-matter pathways through assessment of parameters of water diffusion such as fractional anisotropy (FA), a measure sensitive to degree of myelination [28, 29]. The few human studies assessing associations between white-matter FA and EEG/MEG evoked responses have found that more myelination (as indicated by higher FA values) is associated with earlier visual, auditory, and motor response latencies [30–37]. For example, studies from our laboratory have shown that a more myelinated auditory radiation thalamocortical pathway in school-age children is associated with an earlier latency M50 and M100 evoked response [34–36].

Although marked change in evoked latencies and white matter occur during the first months to the first year of life [17, 18, 26, 27, 38–41], to our knowledge few studies have examined the relationship between evoked response latency and diffusion magnetic resonance imaging (dMRI) measures in infants [visual: 33, auditory: 42]. Adibpour et al. [42] assessed the relationship between transverse diffusivity (a measure of white-matter integrity) and the auditory EEG P2 response to speech sounds in 16 infants (11 males) 1 to 6 months of age. Although an expected relationship between P2 and auditory radiation transverse diffusivity was not significant in this small sample after controlling for age, a trending relationship was observed between left ipsilateral P2 response latency and transverse diffusivity of the auditory callosal fibers [42].

The present study sought to build on this line of research, here examining associations between auditory encoding responses and auditory radiation white matter from birth to 2 years of age. The maturation of auditory cortex neural activity has been studied using EEG [20–25, 43] and MEG [for MEG review see 44]. In infants and young children, the auditory evoked response has a positive-to-negative waveform morphology, with the EEG P2 [also referred to as the P50; 20-22, 24, 45, 46-48] and its MEG equivalent the P2m [also referred to as the P1m or the M50, 25, 49, 50-53] the largest and most consistently observed auditory response in infants [17, 48].

Infant and toddler EEG studies report that P2 latency decreases across young child development [20, 21, 24, 47, 54], with Barnet [20] reporting a 75 ms latency decrease between 10 days and 37 months in a cross-sectional sample. Longitudinal infant EEG studies have reported similar P2 latency changes [21, 22, 24, 47]. MEG studies examining the maturation of the auditory response using adult MEG systems with infants [54–57] and children [35, 39, 57–59] as well as dedicated infant and toddler MEG systems [25, 49, 51–53] have provided further evidence that P2m latency decreases with age [35, 39, 52, 54, 57–59]. In a paper reporting MEG only results in a large sample (overlapping 36% with the present sample), non-linear maturation of the P2m latency was observed, with P2m latencies decreasing rapidly as a function of age during the first year of life, followed by slower changes between 12 and 24 months [17]. Auditory latency changes continue into at least early adolescence, with the P2/P2m response likely developing into the adult P50/M50 response [44], and with an adult-like P50 latency not observed until 12 to 14 years of age [60].

Age-related increases in white matter have been well documented in infant and young child studies [61–68] and may provide a mechanism accounting for age-related reductions in evoked response latency. These studies show that myelination maturation trajectories are regionally specific, with early maturation in the cerebellum, pons, internal capsule, parts of the corpus callosum, and the optic radiations. This is followed by myelination of the occipital and parietal lobes, with temporal and frontal lobe myelination occurring last [for review see 69]. With respect to the auditory cortex, the myelination of auditory radiation white-matter tracts (the thalamocortical tracts sending auditory information from the thalamus to primary/secondary auditory cortex) continues throughout at least the first 3 years of life [3].

### Study Aims

The present study tested the hypothesis that maturation of auditory radiation white-matter microstructure contributes to earlier cortical auditory responses in a cross-sectional cohort of infants 2 to 24 months old. In a smaller sub-cohort with repeated measures, the maturational trajectory of auditory brain development within individual infants was also examined, providing a stronger intra-individual test of the hypothesis. An infant-MEG system developed in our laboratory [59] provided measures of the infants’ auditory cortex neural activity [44, 70], and dMRI provided auditory radiation FA measures. The present study examined the auditory P2m response, generated in left and right Heschl’s gyri [57, 71]. First, auditory cortex function (P2m latency) and auditory radiation white-matter microstructure (e.g., auditory radiation FA) maturation as a function of age for each hemisphere were assessed. Next, associations between auditory radiation FA and P2m latency were assessed. It was hypothesized that significant age-associated changes in P2m latency and auditory radiation FA would be observed and that significant and strong associations between P2m and auditory radiation white matter would be detected, observed even after removing the effect of age on both brain measures.

## Methods

### Participants

This study was approved by the local Institutional Review Board, and all families gave written informed consent. Study data were obtained from an ongoing longitudinal infant multimodal neuroimaging study (R01 HD093776).

All infants were typically developing. Inclusion criteria included (1) no first-degree relative with autism spectrum disorder (ASD); (2) no history of seizure disorder and no first-degree relative with a seizure disorder; (3) no premature birth (< 37 weeks gestation); (4) no non-removable metal in the body; (5) no known hearing or visual impairment (as indicated by passing newborn hearing and vision screening); and (6) no concerns regarding developmental delay (based on parent report and medical records).

The cohort included 95 infants, with their first visit occurring between 2 and 7 months old and with follow-up visits approximately every 5 months. Due to COVID-19, most study families missed one or more visits. In all infants, MEG and MRI data were acquired on the same day. At least one time-point of evaluable MEG data was acquired from 91 infants, and at least one time-point of evaluable MRI data was acquired from 59 infants (the primary limitation being that the infant did not fall asleep for the MRI exam). Of the above, 57 infants had at least 1 time-point of same session MEG + MRI data. Ten of these 57 infants were excluded due to excess movement artifact during imaging (MEG: N = 2; MRI: N = 2) or the infant completing the T1 exam but waking up before completing the dMRI exam (N = 6). Evaluable cross-sectional MEG + structural MRI (sMRI) + dMRI data were thus available from 47 infants 2 to 24 months old (19 females). For the 18 infants with longitudinal data, only a single timepoint was included in the cross-sectional analyses, with in-house code used to randomly select only one time point from each of these infants. Of the 18 infants in the longitudinal cohort (8 females), 16 infants had data from 2 timepoints, and 2 infants from 3 timepoints.

### MEG Auditory Exam

Infant whole-head MEG data were obtained in a magnetically shielded room (Vacuumschmelze GmbH & Co. KG, Hanau, Germany) using an Artemis 123 biomagnetometer (Tristan Technologies Inc., San Diego, CA, USA), with a sampling rate of 5000 Hz and a 0.1 Hz high-pass filter. All infants were scanned when awake in the supine position. The Artemis 123 was designed for use with children from birth to 3 years of age [25, 59]. This system has 123 sensors (first-order axial gradiometers) and a helmet circumference of 50 cm, matching the median 3-year-old male head size in the USA. The Artemis 123 employs a coil-in-vacuum sensor configuration to minimize the distance between the helmet surface and sensors (6 to 9 mm).

Before MEG acquisition, a fabric cap with 4 head position indicator (HPI) coils was placed on the child’s head. The child’s head shape, anatomical landmarks (nasion, right and left preauricular points), and locations of the 4 HPI coils were digitized using a FastSCAN System (Polhemus, Colchester, VT). A research assistant with experience scanning infants and young children stood next to the infant and helped the parent keep the infant calm and alert during the exam. A variety of strategies were used to engage the infant during the scan [see details in 15], including projecting an age-appropriate silent video on a screen placed above the infant’s head (viewing distance 55 cm), providing toys to maintain the infant’s attention and focus, and using a pacifier or bottle. Short breaks were provided as needed. Infants younger than 9 months were swaddled to reduce motion. During the MEG recording, the child’s head position was continuously monitored via the HPI coils.

Auditory stimuli consisted of 500 Hz sinusoidal tones 300 ms in duration. The onset-to-onset interstimulus interval varied between 600 and 2,000 ms. Stimuli were presented via a free-field speaker (Panphonics Sound Shower, Turku, Finland) at 85 dB SPL. MEG recordings were obtained while the infant was awake and alert. The auditory exam lasted 8 minutes (∼300 trials).

### sMRI and dMRI Exams

After the MEG session, sMRI and dMRI data were obtained using a Siemens Prisma 3T MR system with a 32-channel head coil. Before the MRI exam, infants were fed and swaddled. All MRI data were acquired during natural sleep. T1-weighted magnetization-prepared rapid gradient-echo (MP-RAGE) structural images were collected with a field of view of 256 x 256 mm, 208 sagittal slices, and 0.8 x 0.8 x 0.8 mm^3^ spatial resolution. Diffusion MRI (dMRI) data were obtained using a diffusion-weighted spin-echo single-shot EPI sequence with opposed phase encoding direction pairs (AP and PA). Each phase encoding direction acquisition included 7 b=0 volumes, 46 b=1500 s/mm^2^ volumes, and 46 b=3000 s/mm^2^ volumes. Thus, the AP and PA acquisitions each included 99 volumes (98 directions), for a total of 198 volumes. Diffusion sequence parameters were TR = 3222 ms, TE = 89.2 ms, and 1.5 mm isotropic resolution.

Distortion correction based on phase encoding pairs was performed with topup from the FSL analysis package [72, 73]. To correct for artifacts from eddy currents, movements, and intravolume movement, eddy_cuda from the FSL analysis package was run on a graphics processing unit (GPU) cluster [74].

### MEG Source Analyses

MEG data were analyzed using Brainstorm [75] (http://neuroimage.usc.edu/brainstorm). MEG data were downsampled to 1000 Hz and then band-pass filtered from 3 to 55 Hz (transition bands 1.5 to 3.0 Hz and 55 to 63.25 Hz). Heartbeat artifact was removed via independent component analyses (ICA). Other artifacts (e.g., movement, muscle artifact) were visually identified and the effected data were discarded. During the scan, times when the infant was not attending to the stimuli were noted (e.g., crying, falling asleep), and these data segments were manually removed. Other artifacts (e.g., movement, muscle artifact) were visually identified and manually removed. Trials with MEG activity exceeding 500 fT due to excessive motion or magnetic noise artifact, were removed. Averaged head movement across time, recorded from the 4 HPI coils, was computed for each dataset for use as a variable in the statistical analyses.

Auditory evoked responses were obtained by averaging epochs 200 ms pre-stimulus to 500 ms post-stimulus from onset of the stimulus. Across infants, an average of 292 trials were obtained, with an average of 255 ± 60 artifact-free trials.

Digitized surface points from FastSCAN representing the shape of the infant’s head (*>* 10,000 points) were used to co-register each infant’s MEG data to their T1-weighted sMRI using an affine transformation. The T1-weighted data were processed to extract the brain and head surface (needed for source modeling), the surfaces separating the pia mater, gray matter, and white matter estimated using Infant FreeSurfer [107]. The Destrieux [108] cortical parcellations were also computed. Using the auditory evoked response, whole-brain activity maps were computed using Minimum Norm Estimates [MNE; 76, 77-79], available in Brainstorm, with minimum-norm imaging estimating the amplitude of brain sources constrained to the cortex and current dipoles normally oriented to the local surface cortical surface [80]. Auditory evoked-response activity was mapped to each infant’s cortical-surface source space (∼15,000 vertices; varied across infants) as a function of time (one ms resolution). For each infant, an MEG noise covariance matrix was obtained from an empty room recording obtained immediately prior to the infant’s scan. MNE solutions were computed with normalization as part of the inverse routine, based on the noise covariance.

In most infants, the primary/secondary auditory cortex response includes a series of responses of alternating polarity: (1) a peak (N1m) at ∼ 100 ms, followed by (2) a peak (P2m) at ∼ 200 ms, then (3) a peak (N3m) at ∼ 300 ms, and finally (4) a peak (P4m) at ∼ 400 ms. Auditory encoding processes develop rapidly in infants, with P2m peak latency occurring after 200 ms in younger infants and before 200 ms in older infants. Given that the auditory P2m was the strongest and most consistently observed auditory response in our cohort [17, and same observed in 48, 52], analyses focused on the P2m response (see Figure 1 for examples).

**Figure 1.**
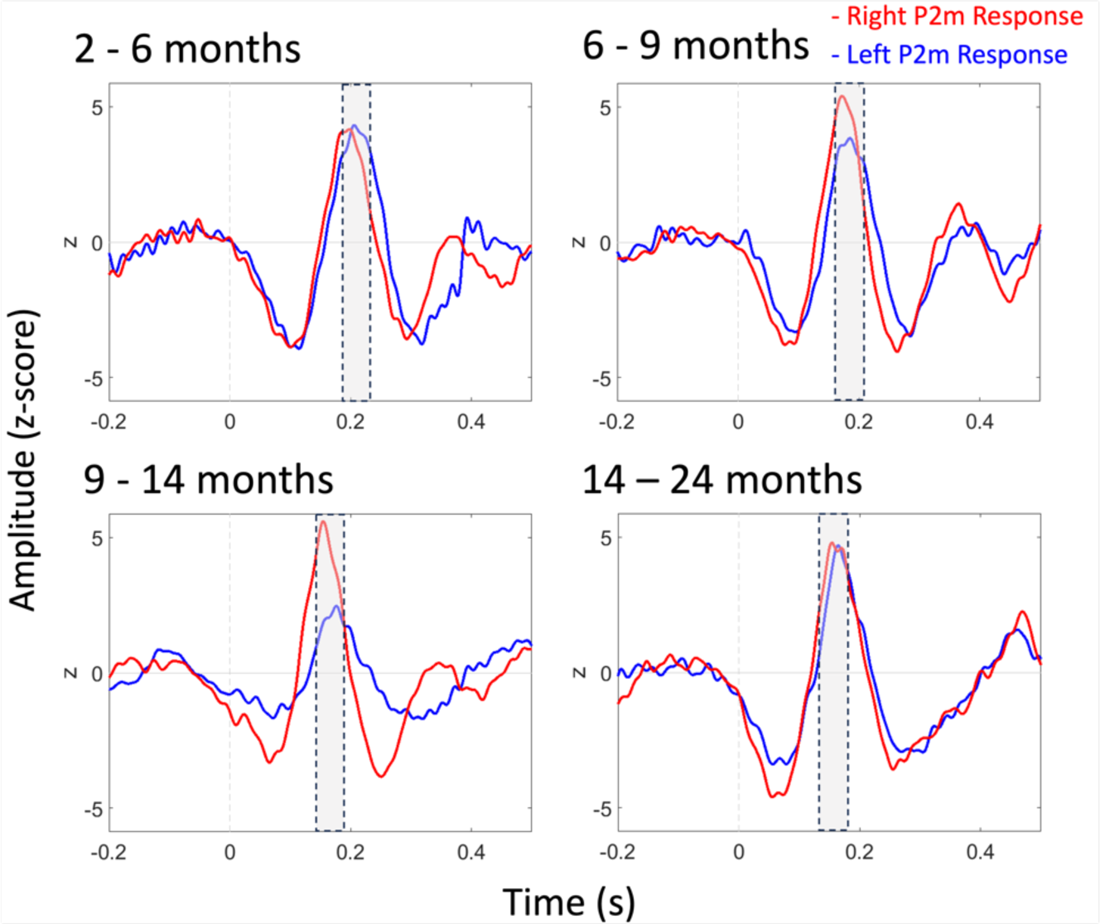
Averaged left (blue) and right (red) auditory source waveforms (from Heschl’s gyrus) in infants 2 to 6 months (N = 11), 6 to 9 months (N = 11), 9 to 14 months (N = 13), and 14 to 24 months (N = 11). The gray bar in each plot highlights the P2m response.

In each infant, left and right hemisphere auditory source timecourses were obtained from left and right hemisphere functional auditory ROIs [81], as well as atlas defined left and right Heschl’s gyrus ROIs defined in the Destrieux Atlas (transverse temporal gyrus). For the functional auditory ROI, the P2m response was first identified via the sensor magnetic field sink and the source topography (e.g., in the left hemisphere an anterior source and posterior sink), and with left and right P2m ROIs created at the time of the peak P2m sensor activity using an amplitude threshold of 5 nAm and a cluster threshold of ≥ 50 neighboring cortical surface vertices. For the functional and atlas-defined ROIs, a left and right auditory source timecourse was obtained via averaging across vertices within the left and right ROI, with post-stimulus activity normalized to the baseline (i.e., providing post-stimulus z-score measures). Using in-house MATLAB software, left and right peak P2m latencies were identified.

### Auditory Radiation White-matter Microstructure

DTI maps were computed using the dtifit function of the FMRIB Diffusion Toolbox (FMRIB Software Library), providing maps of parameters including: fractional anisotropy (FA; degree anisotropy of the diffusion process), mean diffusivity (MD; one third of the tensor trace), axial diffusivity (AD; diffusivity along the main tensor axis), and radial diffusivity (RD; diffusivity perpendicular to the main axis). A study specific infant template was created using Advanced Normalization Tools (ANTs) [82] and each infant’s FA volume was registered to the template. Left and right Heschl’s gyrus regions of interest (ROI) were transformed from template space to each infant’s native space to measure the auditory radiation, the Heschl’s gyrus region capturing the distal portion of the auditory radiations. Auditory radiation measures were obtained via averaging parameters across all voxels in the left and right auditory radiation ROI.

### Statistical Analyses

Statistical analyses were conducted using JMP statistical software (SAS Institute Inc. (2021). JMP Pro, Version 16. Cary, NC: SAS Institute Inc.).

#### Cross-sectional cohort

Associations between age and brain function (P2m) and age and brain structure (auditory radiation white-matter microstructure) were assessed. Given in infants non-linear maturation of the auditory evoked and white-matter measures in infants [109], Akaike information criterion (AIC) were used to determine if a linear or non-linear model best fit the data. Associations between brain structure and function were examined using a mixed model. As an example, a mixed model was run with P2m Latency as the dependent variable, auditory radiation FA, Age (log transformed given a non-linear association between age and the P2m and diffusion measures), Hemisphere, the FA x Hemisphere interaction, and the MEG Head Movement measure entered as fixed effects, and Subject entered as a random effect. Statistical analyses were repeated for auditory radiation MD, RD, and AD. Given significant age-related changes in P2m latency and auditory radiation diffusion measures, to further assess brain function-structure associations, mixed models were run with age-residualized P2m Latency by Hemisphere entered as the dependent variable, age-residualized Diffusion measures and the MEG Head Movement measure entered as fixed effects, and Subject entered as a random effect.

#### Longitudinal cohort

To examine associations between auditory radiation measures and auditory P2m Latency in the children with repeated measures (i.e., follow-up timepoints, see details in Participants section), mixed models were run with P2m Latency entered as the dependent variable, auditory radiation Diffusion measures (separately for FA, MD, RD, and AD), Age (log), Hemisphere, and MEG Head Movement entered as fixed effects, and Subject entered as a random effect.

## Results

### Maturation of P2m Latency and Auditory Radiation White matter

Analyses were conducted for the P2m measures obtained from the left and right functional ROI and the atlas defined Heschl’s gyrus ROI. The pattern of findings was the same using the atlas or functional P2m measures. Given that the auditory radiation white-matter measures were obtained from Heschl’s gyrus, results are only reported using the P2m latencies obtained from the Heschl’s gyrus ROI.

### Cross-sectional cohort

Table 1 shows demographic information and descriptive statistics for P2m latency and auditory radiation diffusion measures for the male and female infants as well as the total sample. Males and females did not differ on age (mean difference 1.9 months, unequal variances *t*(61.35) = −1.68; *p* = .10). T-tests showed males and females did not differ in P2m latency or auditory radiation FA (*ps* > .05). Figure 1 shows averaged left and right auditory evoked responses from Heschl’s gyrus ROIs for infants in 4 different age bins.

**Table 1.** Demographics, descriptive statistics, and the association between Age and P2m latency and auditory radiation measures.

| **Descriptive Statistics** | **Female (N = 19)**<br>**Mean (SD)**<br>**(Min, Max)** | **Male (N = 28)**<br>**Mean (SD)**<br>**(Min, Max)** | **All (N = 47)**<br>**Mean (SD)**<br>**(Min, Max)** | **Nonlinear**<br>**Associations with**<br>**Age: R<sup>2</sup> (p value)** |
| --- | --- | --- | --- | --- |
| **Age (months)** | 10.8 (5.9)<br>(2.9, 22.6) | 8.8 (4.4)<br>(2.2, 18.5) | 9.6 (5.1)<br>(2.2, 22.6) |  |
| **Left P2m Latency (ms)** | 187 (30)<br>(147, 246) | 189 (34)<br>(108, 263) | 188 (32)<br>(108, 263) | 0.28 ($p < .0001$) |
| **Right P2m Latency (ms)** | 178 (36)<br>(118, 250) | 179 (26)<br>(144, 250) | 179 (30)<br>(118, 250) | 0.51 ($p < .0001$) |
| **Left FA (a.u.)** | 0.25 (0.02)<br>(0.22, 0.30) | 0.25 (0.02)<br>(0.22, 0.28) | 0.25 (0.02)<br>(0.22, 0.30) | 0.74 ($p < .0001$) |
| **Right FA (a.u.)** | 0.24 (0.02)<br>(0.21, 0.28) | 0.24 (0.02)<br>(0.20, 0.27) | 0.24 (0.02)<br>0.24 (0.20, 0.28) | 0.72 ($p < .0001$) |
| **Left MD (a.u.)** | 6.8e-4 (4.6 e-5)<br>(6.2e-4, 7.6e-4) | 6.9e-4 (4 e-5)<br>(6.3e-4, 7.6e-4) | 6.8e-4 (4.2 e-5)<br>(6.2e-4, 7.6e-4) | 0.93 ($p < .0001$) |
| **Right MD (a.u.)** | 6.8e-4 (4.1e-5)<br>(6.3e-4, 7.6e-4) | 6.9e-4 (3.2e-5)<br>(6.5e-4, 7.5e-4) | 6.9e-4 (3.5e-5)<br>(6.3e-4, 7.8e-4) | 0.90 ($p < .0001$) |
| **Left RD (a.u.)** | 5.9e-4 (4.8e-5)<br>(5.2e-4, 6.7e-4) | 6e-4 (4.1e-5)<br>(5.4e-4, 6.7e-4) | 5.9e-4 (4.4e-5)<br>(5.2e-4, 6.7e-4) | 0.92 ($p < .0001$) |
| **Right RD (a.u.)** | 6e-4 (4.1e-5)<br>(5.4e-4, 6.9e-4) | 6e-4 (3.3e-5)<br>(5.6e-4, 6.7e-4) | 6e-4 (3.6e-5)<br>(5.4e-4, 6.9e-4) | 0.90 ($p < .0001$) |
| **Left AD (a.u.)** | 8.5e-4 (4.4e-5)<br>(8e-4, 9.4e-4) | 8.7e-4 (3.8e-5)<br>(8.1e-4, 9.4e-4) | 8.6e-4 (4e-5)<br>(8e-4, 9.4e-4) | 0.92 ($p < .0001$) |
| **Right AD (a.u.)** | 8.6e-4 (4e-5)<br>(8e-4, 9.5e-4) | 8.6e-4 (3.1e-5)<br>(8.2e-4, 9.2e-4) | 8.6e-4 (3.5e-5)<br>(8e-4, 9.5e-4) | 0.86 ($p < .0001$) |

When evaluating the maturation of the P2m latency and auditory radiation white-matter measures, a comparison of AIC values showed that for all measures a nonlinear fit was better than the linear fit (see Table 1). Figure 2a shows the non-linear maturation of P2m latency as a function of age in both hemispheres (exponential age transform for left P2m: R^2^ = 0.28, *p* < .0001; exponential age transform for right P2m: R^2^ = 0.51, *p* < .0001), with P2m latencies decreasing rapidly as a function of age during the first year of life, followed by slower changes from 12 to 24 months. Figure 2b shows the non-linear maturation of auditory radiation white-matter microstructure (FA, MD, AD, RD) in both hemispheres, with auditory radiation white-matter measures changing rapidly as a function of age during the first year of life, followed by slower changes from 12 to 29 months (see Table 1 for the summary of fits of all measures). Of note, MEG head movement was not associated with age.

**Figure 2.**
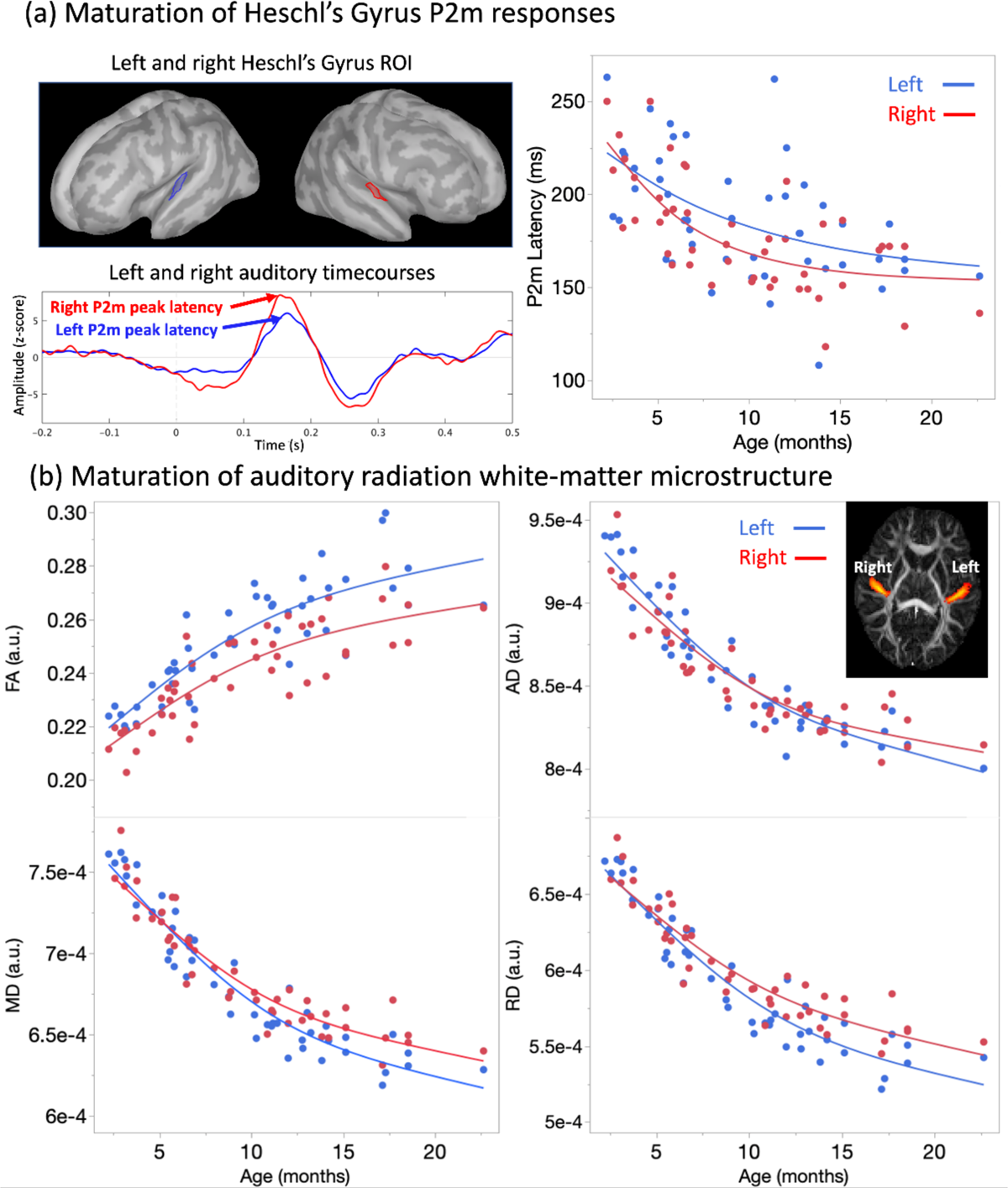
Maturation of P2m latency and auditory radiation FA. (a) Left panel: Heschl’s gyrus P2m neural activity (top panel) and the associated left (blue) and right (red) auditory cortex source waveforms (bottom panel) in a 5-month-old infant. Right panel: A non-linear age-related decrease in P2m latency was observed bilaterally. (b) Non-linear age-related increase in auditory radiation FA, MD, AD, and RD in both hemispheres.

### P2m Latency and Auditory Radiation White-matter Associations

#### Cross-sectional cohort

Table 2a shows summarizes the mixed model results, performed with P2m Latency as the dependent measure, auditory radiation Diffusion (FA, MD, RD, and AD entered as predictors in separate mixed models), Age (log), Hemisphere, and Head Movement as fixed effects, and Subject as a random effect. The interaction term for Hemisphere as well as the 4 diffusion measures were not significant and were thus removed from the mixed models so that the main effects could be directly interpreted. As shown in Table 2a, Hemisphere as well as most of auditory radiation diffusion measures (see Figure 3) accounted for significant variance in P2m latency. The Hemisphere main effect showed earlier P2m responses in the right than left hemisphere (see Table 1 and Figure 2a). The Figure 3a scatterplots show the auditory radiation diffusion measures and P2m latency association in each hemisphere, with these associations well represented by a linear fit. For example, the left auditory radiation MD accounted for 34% of the variance in left P2m latency, and right auditory radiation MD accounted for 50% of the variance in right P2m latency.

**Figure 3.**
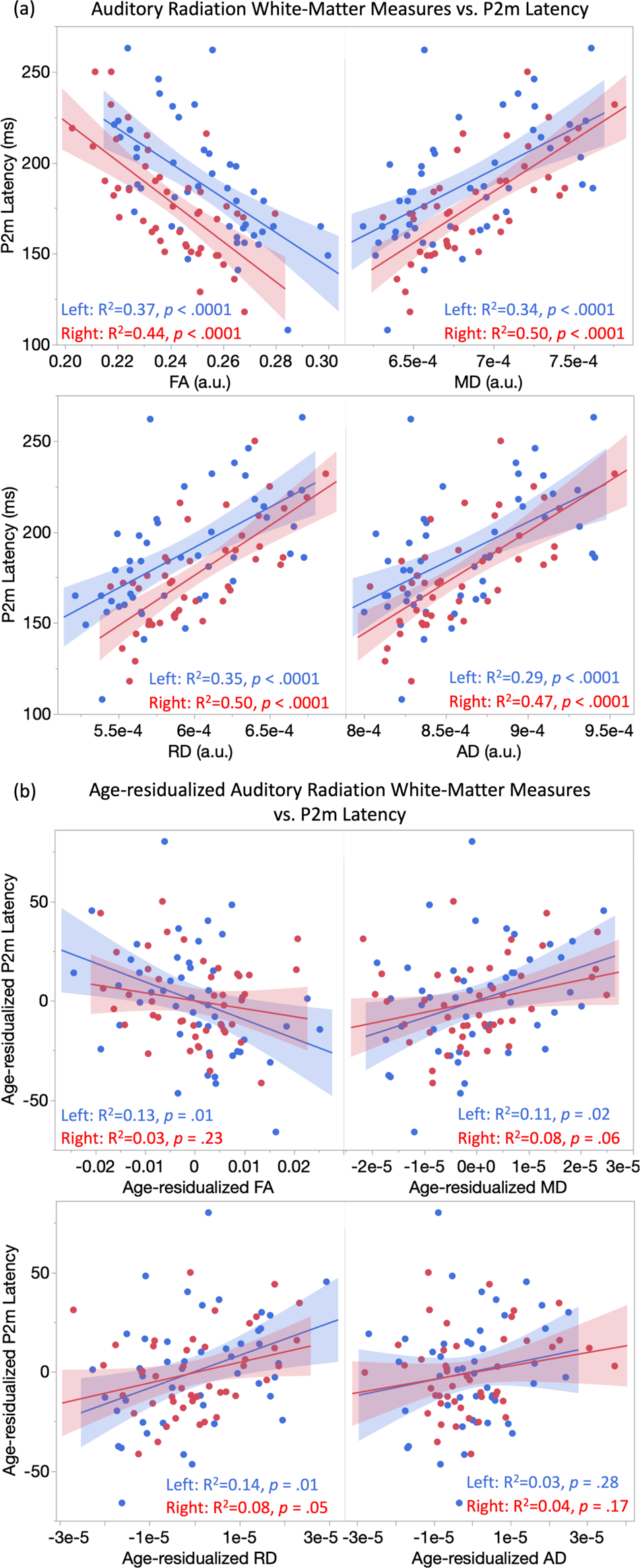
(a) Associations between auditory radiation diffusion (x axis) and P2m latency (y axis), with the 95% confidence interval (shading) for the left hemisphere (blue) and right hemisphere (red). (b) Associations between age-residualized auditory radiation diffusion (x axis) and age-residualized P2m latency (y axis).

**Table 2a.** Summary of linear mixed model results for the cross-sectional cohort.

| Predictors | FA |  | MD |  | RD |  | AD |  |
| --- | --- | --- | --- | --- | --- | --- | --- | --- |
|  | F Ratio | *p* | F Ratio | *p* | F Ratio | *p* | F Ratio | *p* |
| Auditory Radiation | 5.00 | 0.03 | 4.77 | 0.03 | 6.17 | 0.01 | 0.93 | 0.34 |
| Age (log) | 2.90 | 0.09 | 0.00 | 0.99 | 0.02 | 0.88 | 1.85 | 0.18 |
| Hemisphere | 11.28 | < 0.01 | 9.58 | < 0.01 | 11.44 | < 0.01 | 5.96 | 0.02 |
| Head Movement | 0.18 | 0.67 | 0.06 | 0.81 | 0.09 | 0.76 | 0.06 | 0.81 |

Given significant age-related changes in P2m latency and auditory radiation diffusion measures (see Figure 2), to further assess brain function-structure associations, follow-up mixed models were run with age-residualized P2m Latency the dependent variable, Head Movement and each age-residualized Diffusion measure as fixed effects, and Subject a random effect. As shown in Table 2b, the diffusion and P2m findings remained significant. As an example, after regressing out age from P2m latency and auditory radiation MD, age-residualized auditory radiation MD still accounted for significant variance in age-residualized P2m latency (*p* = .01). Of note, among all diffusion measures, auditory radiation AD does not account for significant variance in P2m latency, with or without removing age in the brain function and structure measures. The Figure 3b scatterplots show that even in this very conservative condition the associations between the age-residualized auditory radiation diffusion measures and age-residualized P2m latency remained significant even after removing the variance associated with age from both brain measures. As an example, after regressing out age from both brain measures, the left auditory radiation MD accounted for 11% of the variance in left P2m latency, and right auditory radiation MD accounted for 8% of the variance in right P2m latency.

**Table 2b.** Summary of linear mixed model results for cross-sectional cohort after removing variance in the brain measures associated with age.

| Predictors | Age-residualized FA |  | Age-residualized MD |  | Age-residualized RD |  | Age-residualized AD |  |
| --- | --- | --- | --- | --- | --- | --- | --- | --- |
|  | F Ratio | *p* | F Ratio | *p* | F Ratio | *p* | F Ratio | *p* |
| Age-residualized Auditory Radiation | 6.40 | 0.01 | 6.29 | 0.01 | 8.00 | < 0.01 | 1.19 | 0.28 |
| Head Movement | 0.30 | 0.58 | 0.13 | 0.72 | 0.18 | 0.67 | 0.13 | 0.73 |

#### Longitudinal cohort

The relationship between P2m Latency and auditory radiation Diffusion was evaluated in the 18 infants with multiple time-point data. As shown in Table 3, results showed main effects of Diffusion and Hemisphere (left P2m latency > right P2m latency), with each of the auditory radiation diffusion measures except for AD associated with P2m latency. Figure 4 shows left and right auditory radiation FA and P2m latency trajectories for each infant.

**Figure 4.**
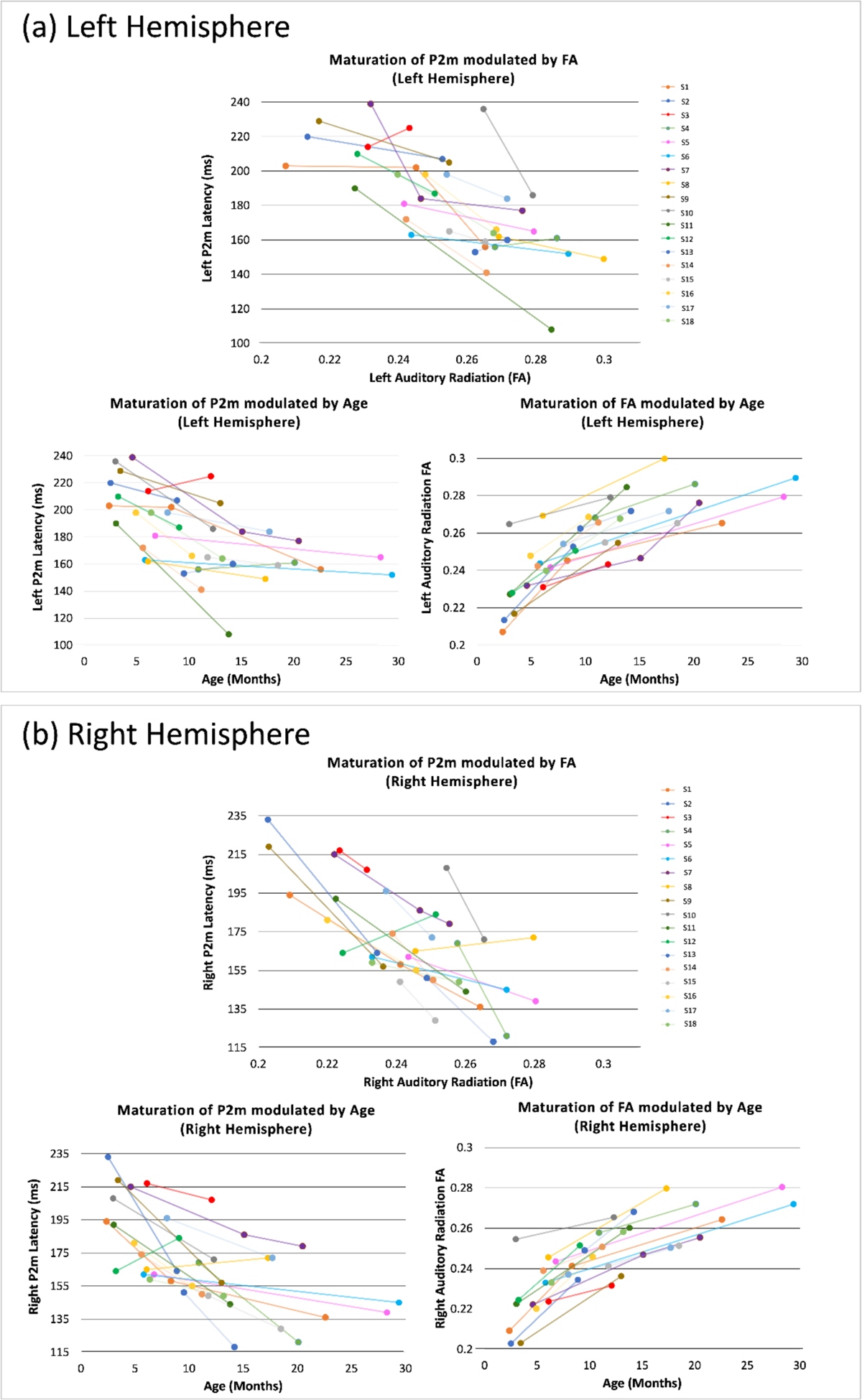
Maturational trajectory of P2m latency and auditory radiation FA. Each colored line represents a single infant. (a) Top panel: Associations between left auditory radiation FA (x axis) and left P2m latency (y axis) across time; Bottom left panel: Associations between age (x axis) and left P2m latency (y axis) for each infant; Bottom right panel: Associations between age (x axis) and left auditory radiation FA (y axis) for each infant. (b) Same for right hemisphere.

**Table 3.** Summary of linear mixed model results for the longitudinal cohort.

| Predictors | FA |  | MD |  | RD |  | AD |  |
| --- | --- | --- | --- | --- | --- | --- | --- | --- |
|  | F Ratio | *p* | F Ratio | *p* | F Ratio | *p* | F Ratio | *p* |
| **Auditory Radiation** | 12.99 | < 0.01 | 12.17 | < 0.01 | 17.65 | < 0.01 | 3.64 | 0.06 |
| **Age (log)** | 0.04 | 0.85 | 2.30 | 0.13 | 4.68 | 0.03 | 0.14 | 0.71 |
| **Hemisphere** | 25.81 | < 0.01 | 22.16 | < 0.01 | 28.15 | < 0.01 | 12.71 | < 0.01 |
| **Head Movement** | 0.02 | 0.88 | 0.05 | 0.83 | 0.02 | 0.88 | 0.04 | 0.84 |

## Discussion

The present study provides strong support for the hypothesis that an increase in thalamocortical neural conduction velocity, due to increased axon diameter and/or myelin maturation, contributes to a decrease in the infant P2m auditory evoked response latency. Both the cross-sectional and longitudinal data provided statistical support for this association. Whereas the cross-sectional model showed an association between brain structure and function across a group of infants even after removing effects of age from both brain measures, the longitudinal model showed that acoustic radiation diffusion and P2m latency associations could be observed at the level of a single child.

### Maturation of Auditory Cortex in Infants and Toddlers

Consistent with our recent study reporting cross-sectional auditory P2m findings in a larger cohort with 114 infants [81], a nonlinear age-associated decline in left and right P2m latency was observed, with a more rapid change in P2m latency during the first than second year of life (see Figure 2a). Hemisphere P2m latency differences were observed, with an earlier P2m latency in the right than left hemisphere, this lateralization finding consistent with that observed in other infant [81] [83–87] and older child auditory studies [25, 39, 52, 57, 60, 88].

A non-linear age-associated change in the left and right auditory radiation diffusion measures was also observed, with the finding of a more rapid change in auditory radiation white-matter microstructure during the first than second year of life consistent with prior studies [106].

### Association between Auditory Cortex Function and Structure

As noted in the Introduction, understanding the relationship between functional and structural development of the auditory system in infants is important as rapidly encoding auditory information is necessary for parsing speech sounds and learning to map meaning onto sounds [6–11]. The present study shows an association between auditory P2m latency and auditory radiation diffusion measures in children 2 to 29 months old, with these findings parsimoniously interpreted as showing that increased axon diameter and/or myelin maturation in the auditory radiation thalamocortical tract fosters faster transmission of auditory information from the thalamus (primarily the medial geniculate body) to the primary/secondary auditory cortex. These results are consistent with our studies exploring associations between auditory neural activity and local white matter in older children [34–36].

Present results differ from an infant study exploring the relationship between the auditory EEG P2 latency and white matter. Adibpour et al. [42] did not find an association between the auditory P2 latency and auditory radiation transverse diffusivity in 16 infants 1 to 6 months old when controlling for age. In the present study, left and right auditory radiation diffusion measures predicted from 29% to 50% of the variance in P2m latency (see Figure 3). Different from Adibpour et al., the present study showed that the auditory radiation RD (a measure of water movement across fibers; the same measure used in Adibpour et al.) predicts variance in P2m latency. A difference in findings between Adibpour et al. study and the present study may be due to a smaller sample in Adibpour et al. not providing the power needed to detect associations, as well as obtaining their auditory latency measures using scalp EEG, with simulations as well as empirical work indicating that left and right auditory cortex latencies may be difficult to estimate using EEG given the orientation of auditory cortex neural generators [89, 90].

In the present study, all diffusion measures (except auditory radiation AD) predicted variance in P2m latency. For example, left and right auditory radiation MD predicted a significant 34% (left) and 50% (right) of the variance in left and right P2m latency. When using the age-regressed brain function and structure variables, age-residualized auditory radiation MD accounted for 11% and 8% of the variance in age-residualized left and right P2m latency, respectively, this considered a strong effect [91]. Age is a surrogate for multiple biological factors which modulate P2m latency, with white-matter microstructure as well as many other biological factors contributing to rapid conduction velocity. The age-residualized association measures are a conservative low-end estimate, as it is probable that removing age from both brain measures removes real shared variance between the auditory radiation diffusion measures and P2m latency.

### Limitations

The present study focused on associations between primary/secondary cortex neural activity and the associated thalamocortical pathways with these measures both functionally and geographically related through their structural connectivity. Previous studies have found associations between more geographically distant sources of neural activity and white matter. For example, Adibpour [42] reported a negative trending relationship between the left ipsilateral EEG P2 auditory response and transverse diffusivity of the auditory callosal fibers, and Stufflebeam et al. [30] reported a negative association between the peak latency of the occipitally recorded visual evoked response with bilateral parietal and right frontal FA. As our infant sample size increases, future analyses will examine associations between left and right P2m latency and white-matter measures throughout the brain. Finally, although sex differences and hemisphere differences were not observed, the present study was underpowered to detect what are likely small to medium effects.

## Conclusions

The present infant study provides strong support for the hypothesis that maturation of thalamocortical white matter and myelination contributes to an increase in neuronal conduction velocity and a resultant decrease in auditory evoked response latency. An understanding of how brain function and structure in the auditory system are associated in typically developing infants and children is needed to better understand and identify differences in brain development in children with clinical disorders, especially clinical disorders showing differences in auditory encoding neural activity.

## Acknowledgements

The authors would like to thank the subjects who participated in the studies and John Dell, Rachel Golembski, Erin Huppman, Peter Lam, Shivani Patel, Na’Kiesha Robinson, Michelle Slinger, and Taylor Chiang who helped with data collection. The authors also thank Prof. Gregory A. Miller for comments on an early draft of the paper.

## Funding

This work was supported by the National Institute of Child Health and Human Development (NICHD) (R01HD093776 to Dr. J. Christopher Edgar; P50HD105354 to Drs. Roberts and Edgar); the National Institute of Mental Health (NIMH) (R01MH107506 to Dr. J. Christopher Edgar; K01MH108822 to Dr. Yuhan Chen); the Eagles Autism Foundation (Pilot Grant to Dr. Yuhan Chen); and the Nancy Lurie Marks Family Foundation to Dr. Heather L. Green (principal investigator Roseann Schaff).

## Author Contributions

YC, TPLR, and JCE designed the study. YC, HLG, MP, and JCE conducted the statistical analyses. YC, HLG, JB, SL, and JCE developed and implemented data acquisition and/or processing pipelines. ESK and MK developed the clinical and cognitive assessment battery. YC, HLG, JB, MP, TPLR, and JCE made substantial contributions to the interpretation of the data. YC, HLG, KM, MM, OA, TY, and MK participated in data acquisition and analysis. YC, HLG, and JCE drafted the manuscript. All authors substantively contributed. All authors read and approved the final manuscript.

## Notes

### Competing Interest Statement

The authors have declared no competing interest.

